# An Entrainment Oscillator Mechanism underlies Human Beat Matching Performance

**DOI:** 10.1101/2024.03.07.583955

**Authors:** Chloe Mondok, Martin Wiener

## Abstract

Humans possess an innate ability to effortlessly entrain to auditory rhythms, which previous theories have linked to the supplementary motor area (SMA). Yet, whether entrainment, as measured by electrophysiological (EEG) recordings reflects actual processing of rhythms or merely a reflection of their periodical nature, is unknown. Here we conducted tested human participants on a novel beat matching task, in which they listened to two simultaneous tempos and asked to modulate the rate of a variable tempo (1.67-2.34 Hz between trials) to match a constant target tempo (2 Hz). EEG recordings exhibited entrainment to both frequencies at frontocentral electrodes that shifted into alignment over the course of each trial Behaviorally, participants tended to anchor the matched tempo to the starting comparison frequency, such that they underestimated the tempo for slower initial conditions and overestimated for faster initial conditions; further, tempo judgments were shifted away from the variable tempo on the previous trial. A model of phase-coupled oscillators, in which both tempos were pulled towards one another, replicated both effects. This model further predicted that by strengthening the coupling strength of the constant tempo oscillator, both bias effects could be eliminated. To test this, a second group of subjects underwent transcranial alternating current stimulation (tACS) to the SMA phase-locked to the target tempo at 2 Hz. Consistent with model predictions, tACS attenuated both behavioral effects. Overall, these results provide causal support to the role of the SMA in entrainment during human beat matching.

## Introduction

Humans have the unique intrinsic ability to entrain to auditory stimuli. This is most noticeable through physical synchronization to musical beat but is also present at the neural level. A prominent feature of neural entrainment is coupled neural firing to external auditory stimuli in which both oscillate synchronously (Ross & Balasubramaniam, 2022). In this way, neural populations can sync to external stimuli such that phase and firing frequency are locked to the exogenous stimuli (Haegens & Zion Golumbic, 2018; Lakatos et al., 2019; Ross & Balasubramaniam, 2022; Rosso et al., 2023). Whether specific regions of the brain are eliciting true endogenous neural entrainment or rather mimicking exogenous stimuli internally is not well understood in the literature. That being said, rhythm provides a potential avenue for investigating neural entrainment.

Rhythm is defined as a systematic arrangement of a regularly repeated pattern of sounds known as beats. Though it may not seem necessary for beat perception at first, the integration of the motor system allows for improvements in beat perception (Cannon & Patel, 2021). In fact, the motor system has been found to be capable of mapping out beat intervals themselves (Cannon & Patel, 2021). Further, even in the absence of movement, areas that are a part of the direct pathway of the motor network are active, most notably the supplementary motor area (SMA) and striatum (Cannon & Patel, 2021).

The supplementary motor area (SMA), located in the dorsomedial prefrontal cortex, has recently gained attention in the scientific community for its elusive role in time perception (Clairis & Lopez-Persem, 2023). Indeed, prominent theories of time perception include the SMA (Behrens et al., 2013; Naghibi et al., 2022; Schwartze et al., 2012; Wiener et al., 2010). The leading theory consists of a distributed network of cortical and subcortical areas that are specialized for timing conditions involving core regions such as the cerebellum, SMA, premotor cortex, basal ganglia (Allman et al., 2014; Cona & Semenza, 2017; Coull et al., 2015; Mioni et al., 2020; Nachev et al., 2008; Teki et al., 2012). A key feature of the SMA is that it is involved in both motor and non-motor tasks and, in particular, seems to create a bridge between the two networks. As such, the integration of sensorimotor systems with a forward prediction model allows for a stabilization of the internal timing interval, which is essential for beat perception in music (Amrani and Golumbic, 2022; Rosso et al., 2023).

Given its involvement in both the motor and auditory pathways, the SMA has been shown to play a role in beat perception, the ability to recognize a steady beat due to rhythm (Cannon & Patel, 2021). This is most likely due to its sequencing capabilities in which the SMA can track timing intervals through the use of a forward prediction model. According to the action simulation for auditory prediction (ASAP) hypothesis, information about rhythm is sent from the auditory cortex to motor planning areas where an artificial action for the interval of the beat is created and sent back to the auditory cortex in alignment with the next anticipated beat (Cannon & Patel, 2021). This exchange of information between cortices repeats for each perceived beat, allowing for continuous updates in real time. It is believed that the SMA is responsible for this beat-based action planning. In line with this, Grahn and Rowe (2012) found that the SMA had more activity for consistent rhythms that maintain the beat compared to irregular rhythms with nonbeats. Additionally, it was found that the SMA fired in time to the tempo (Caden-Valencia, et al. 2018), along with other areas such as auditory and sensorimotor cortices resulting in cortico-cortical coherence in which numerous regions across the cortex fire in tandem to the external stimuli (Fujioka et al., 2012; Ross and Balasubramaniam, 2022).

As such, though the SMA is active during overt inter-beat interval timing tasks, recent research also shows that it is active in covert beat perception. Cannon and Patel (2021) suggest that the SMA tags intervals via dynamic firing rates allowing for relative timing information by tracking the stimulus phase regardless of the presence of overt movement (Rosso et al., 2023). Indeed, this dynamic state is believed to be an efference copy of the motor movements, allowing for a forward prediction model that works in tandem with sensory feedback, which is used for beat perception (Cannon & Patel, 2021). In this way, the SMA acts as an internal metronome by replicating the beat at the neuronal level, allowing people to process rhythm when listening to music (Cadena-Valencia et al., 2018; Cannon & Patel, 2021). The idea of the SMA acting as an internal metronome was demonstrated by Cadena-Valencia and colleagues (2018). Their paradigm consisted of training rhesus monkeys to continue the intervals of a visual metronome after it had been removed. They found that the SMA continued to fire gamma bursts at the interval of the metronome in its absence demonstrating that the SMA maintained the beat internally even after the external stimuli was removed (Cadena-Valencia et al., 2018). This is further supported in research by Iverson and colleagues (2009) who found that beta band oscillations (20-30 Hz) resembled that of externally metered beats when participants were asked to intrinsically apply meter to a rhythmically ambiguous phrase. As such, even though the participants were not hearing a meter, when asked to apply an endogenous meter, the SMA acted similarly to that of the exogenous meter (Iverson et al., 2009; Ross and Balasubramaniam, 2022). These support the role of the SMA in beat perception regardless of the inclusion of motor movement.

The SMA’s involvement with the motor system, particularly motor planning, is believed to be responsible for its predictive capabilities when it comes to time perception. A key component of forward prediction modelling for time perception is adjusting predictions in response to error detection. As such, the SMA can compensate for these errors by correcting its timing interval based on feedback. The exact nature of the SMA’s role in error detection is unknown though present. Ide and Li (2011) found that the SMA was activated along with the ventrolateral prefrontal cortex (VLPFC) during behavioral changes due to error correction. Similarly, the SMA is active during target detection in oddball paradigms, further supporting the idea that the SMA can detect errors through monitoring (Linden et al., 1999). This is also seen in musical contexts. When temporal deviations were presented to participants in a passive listening paradigm in which interonset intervals deviated from the established tempo, larger ERP/ITPC were elicited in the frontocentral region (Menceloglu et al., 2020). This effect is seen for both shorter and longer interonset intervals (Ford & Hillyard, 1981; Menceloglu et al., 2020).

Though we know that the SMA is involved with beat perception, the question still remains on its exact role in this process. Given that the SMA utilizes forward prediction models, do levels of activation change between error detection and error correction in rhythm? Nachev and colleagues (2008) discuss how activation of the SMA, specifically the pre-SMA, is seen in changing and inhibiting eye and hand movements; however, is this translational to beat perception when aspects of the beat, rather than body movements, change? Specifically, is there a change in SMA activity for purely perceptual conditions for beat perception in the absence of motor movements? A growing area of interest as to why this may be is the SMA’s ability to entrain, in which neural oscillations couple to exogenous rhythmic stimuli. As such, the SMA is able to modulate its neural firing to sync to the external beat in order to improve temporal predictions of rhythmic intervals (Cadena-Valencia et al., 2018; Cannon & Patel, 2021). That being said, whether neural entrainment is actually occurring or whether the brain is mimicking exogenous auditory stimuli endogenously is still debated.

To address this, we developed a novel tempo matching task in which participants were presented with two simultaneous sounds (a bass tone and a musical tone) at different tempos and asked to match the tempos by adjusting that of the musical one using a keyboard (Figure 1, see Methods). Each trial was self-modulated by the participant and was finished when the participant believed the tempos matched. This task was used across two experiments: Experiment 1 (which employed EEG) and Experiment 2 (which employed tACS which was set to phase-lock the target tempo at 2 Hz for each trial).

**Figure 1:**
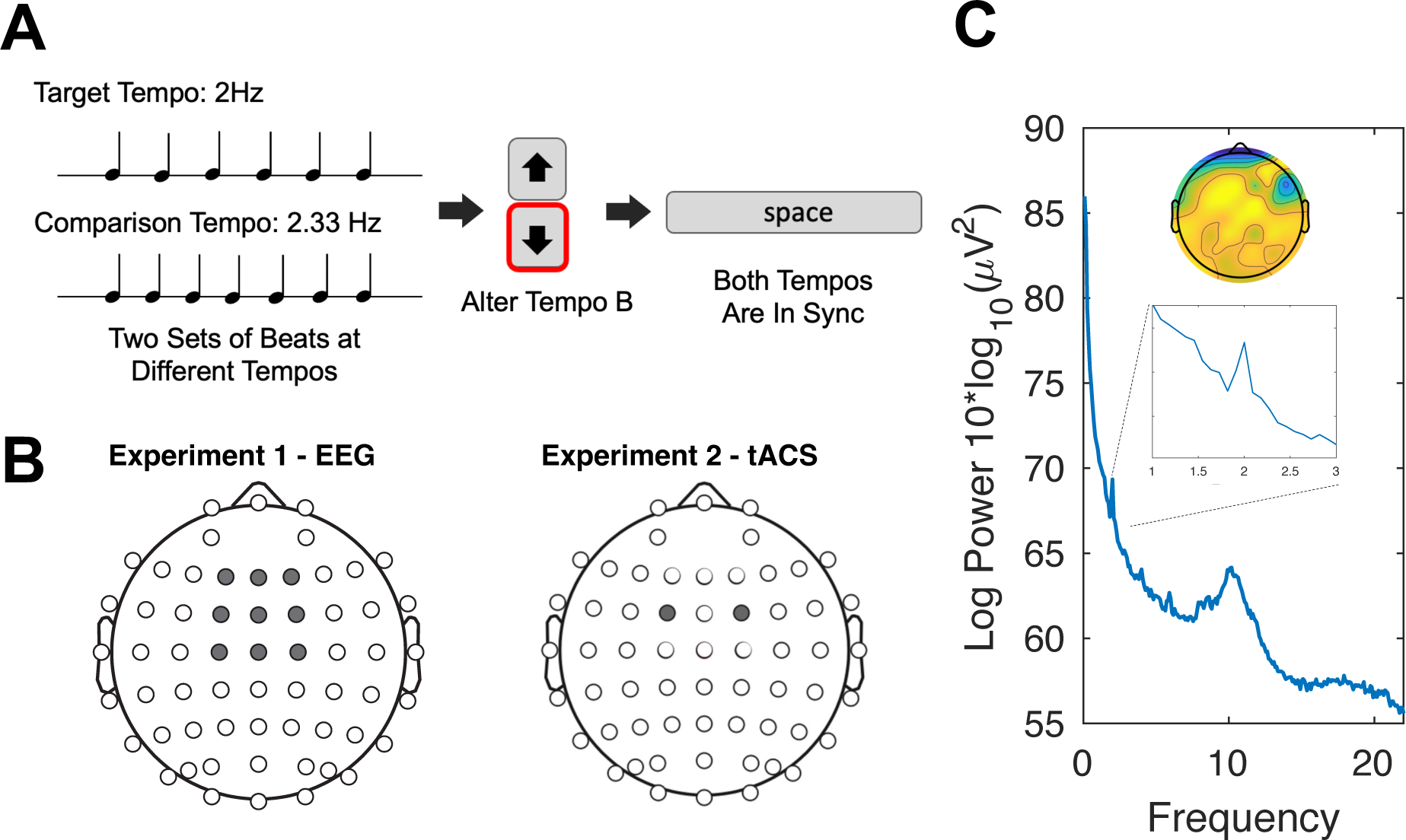
Task Design. **A)** Participants performed a beat-matching task in which they were presented simultaneously with two auditory tempos: a standard 2Hz stimulus and a comparison stimulus with variable frequency between trials (1.66-2.33Hz). Subjects altered the rate of the variable tempo using up and down keyboard presses until they judged them as synchronous, at which point a response was submitted. **B)** Electrode montages used for Experiments 1 and 2. In Experiment 1, 64-channel EEG were collected, with a frontocentral cluster of interest, centered on electrode FCz. In Experiment 2, tACS was applied via two electrodes placed at FC1 and FC2. **C)** Average EEG frequency spectrum from baseline trials of Experiment 1 (2Hz tempo only), demonstrating a peak at the beat frequency (inset). Scalp map displays the average scalp distribution at 2Hz, with maximal power at frontocentral electrodes.

## Results

### Experiment 1

#### Behavioral Data

Participants’ final comparison frequency from each trial was averaged across each condition. We ran a repeated measures ANOVA and found a significant effect for comparison frequencies of that trial (*F*_(5,110)_ = 32.35, *p* < .001, 17^2^*_p_* = 0.60; Fig. 2B), in which participants tended to anchor their matched frequency to that of the comparison frequency in a linear fashion (*t*_(110)_ = 12.38, *p* < .001). For trials that presented comparison frequencies that were lower (1.66, 1.75, or 1.83 Hz) than the target frequency (2 Hz), participants tended to have lower corrected frequencies (*M* = 1.86) while when presented with the higher comparison frequencies (2.16, 2.25, or 2.33 Hz) than the target, participants tended to have higher corrected frequencies (*M* = 2.03). Though lower comparison frequencies tended to be underestimated while higher comparison frequencies tended to be overestimated, on average, higher comparison frequencies were corrected closer to the target frequency than the lower comparison frequencies.

**Figure 2:**
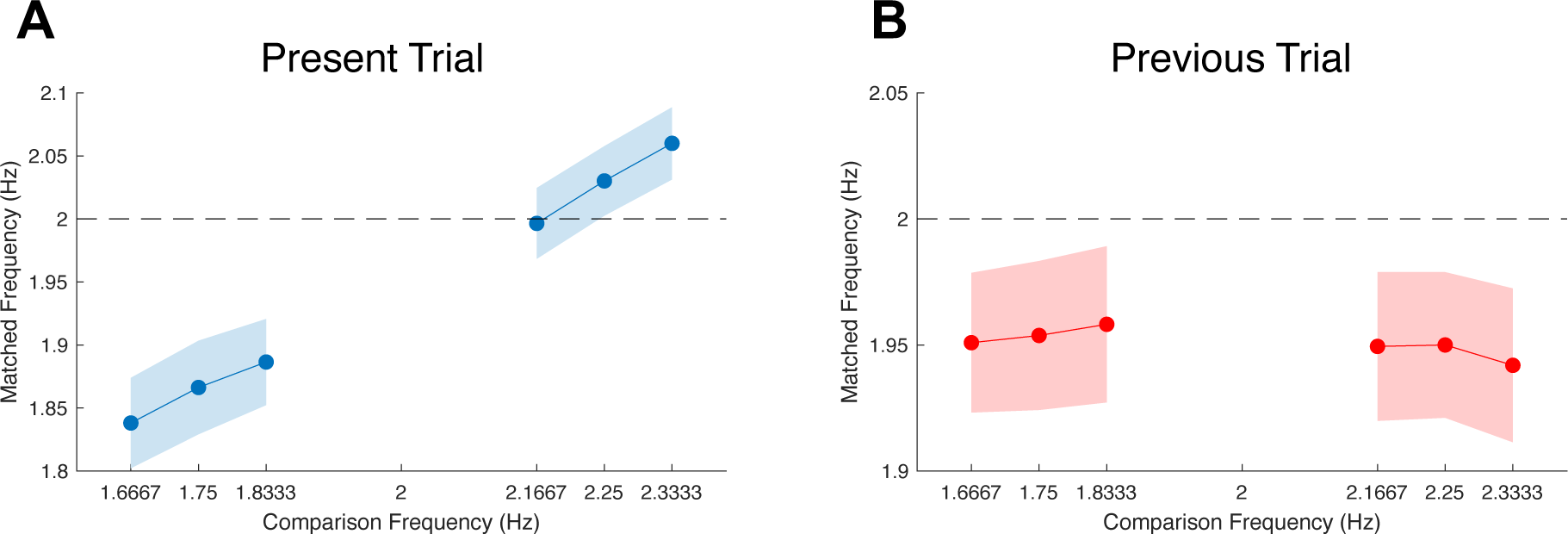
Behavioral data from Experiment 1. A) Average reported frequency of each comparison frequency, demonstrating a bias in synchrony judgments such that comparison frequencies that start at faster/slower rates are biased in the direction of those rates (dashed line represents the target tempo frequency and where correct performance would lie). (B) Average reported frequency by comparison frequency presented on the previous trial, in which matched frequencies were faster/slower rates on the previous trial led to slower/faster judgments on the present one. Shaded regions represent between-subject standard error.

Why might subjects exhibit this bias? Notably, there is no reason to anchor the corrected frequency, as participants have total control over the variable tempo and can easily shift it into alignment regardless of where it starts. We considered that difficulty affected the judgment, yet found no relation between average trial time and the anchoring effect [Spearman’s rho = −0.22, *p*=0.29]. The alternative possibility is that the anchoring reflects a perceptual effect; that is, subjects perceived the stimuli as synchronous even if they objectively were not. If this effect were perceptual, rather than decisional and driven by difficulty, one might expect adaptation-level effects to the continuous presentation of tempo (Becker & Rasmussen, 2007; Wiener, Thompson, & Coslett, 2014). In support of this, a repeated measures ANOVA was run to compare the matched frequency of the current trial to the comparison frequency of the previous trial (n – 1). A significant effect was found for the presentation of the previous trial on the present trial (*F*_(5,100)_ = 2.48, *p* = .036, 17^2^*_p_* = 0.11; Fig. 2B) which had a linear relationship, but in the opposite direction as the present trial data (*t*_(110)_ = −3.05, *p* = .003). As shown in Fig. 2B, responses tended to shift away from the frequency presented on the previous trial, while also in general being slower than 2Hz. This can be seen through the slower conditions having a higher matched frequency for *n* – 1 compared to *n* (Fig. 2) as well as the faster conditions having a lower matched frequency. Further, both behavioral biases were correlated, such that subjects who exhibited a larger bias to the starting comparison frequency also exhibited a larger carryover effect of the previous trial comparison frequency [Spearman rho: −0.45, *p*=0.04].

#### EEG Data

Baseline EEG activity was collected prior to the beginning of the experiment and consisted of participants listening to the bass tone at 2 Hz for 5 minutes. Baseline EEG activity showed a peak at the target frequency (2Hz) in the frontocentral electrodes centered around FCz (Fig. 1C).

Frontocentral EEG activity at the onset of each trial showed peaks at the target frequency and the comparison frequencies (1.66, 1.75, 1.83, 2.16, 2.25, and 2.33 Hz). We ran a repeated measures ANOVA and found that there was a significant effect of frequency per trial onset (*F*_(5,110)_ = 5.59, *p* < .001, 17^2^*_p_* = 0.20; Fig. 2) in a linear fashion (*t*_(110)_ = 4.79, *p* < .001). Indeed, at the beginning of each trial, frontocentral EEG activity signaled both the target frequency as well as the comparison frequency presented in that trial. A significant effect was not found for frequencies for trial offset (*F*_(5,110)_ = 1.92, *p* = .097, 17^2^*_p_* = 0.08; Fig. 2) though a linear relationship was still present (*t*_(110)_ = 2.17, *p* = .032).

#### Oscillator Model

To explain the behavioral and EEG results observed, where participants exhibited attractive bias to the starting frequency of the comparison tempo in synchrony judgments and repulsive bias from the frequency on the previous trial, we adapted a model of coupled oscillators, based on previous work (Zalta, et al. 2020). In this model, two phase-coupled oscillators, corrupted by Gaussian noise and driven by the standard (S) and comparison (C) frequencies, exert a coupling strength (K) on each other across a simulated trial (Fig. 4A). If the coupling strength of the standard is dominant and exclusive, then the comparison oscillator will be pulled into alignment at the same frequency as the standard. However, if the comparison frequency exerts its own coupling strength, then the final comparison frequency will always remain away from the standard frequency in the direction of the comparison. We tested this model on the same number of trials as human participants (*n* = 120); crucially, we also introduced between-trial adaptation in the model, such that the standard was pushed away from the final frequency on the previous trial. Here, we observed that this model replicated both bias effects in behavioral data, such that the final comparison frequency on each trial was linearly shifted in the direction of the starting comparison frequency; further, the comparison frequency shifted in the opposite direction of the previous trial frequency (Fig. 4B).

**Figure 3:**
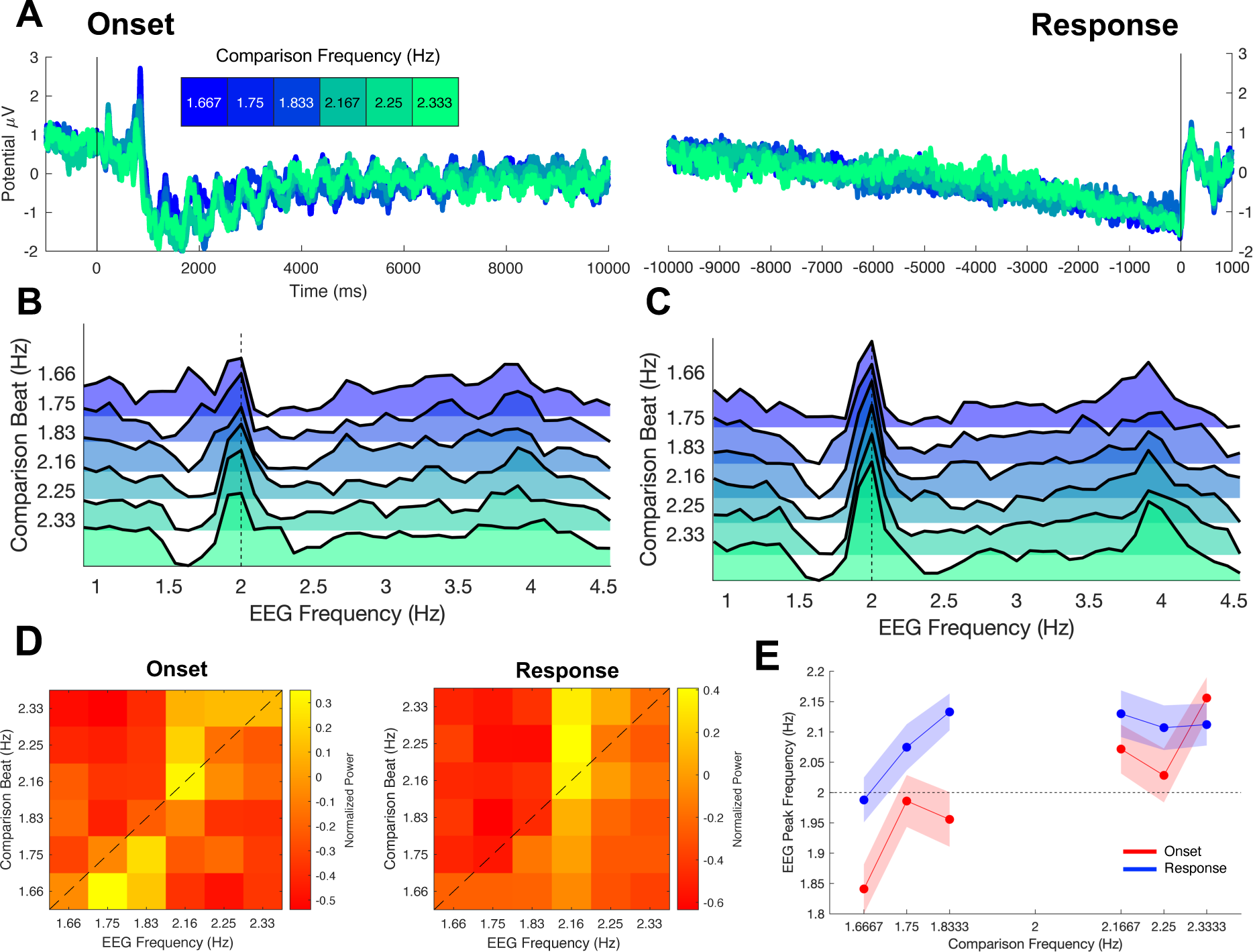
EEG data from Experiment 1. **A)** Grand-average Event-related potentials (ERPs) time-locked to the start (onset) and end (response) of each trial for each comparison frequency. **B**) Normalized frequency spectra for each comparison frequency for onset-locked data demonstrating a peak at 2Hz and a sub-peak at the comparison frequency. **C)** Frequency spectra at response, demonstrating higher peaks at 2Hz, but with smaller sub-peaks at comparison frequencies. **D)** Heatmaps of normalized power at each of the comparison frequencies within the same EEG frequencies; if the comparison frequency exerts an effect, the highest power should be observed along the dashed diagonal line, which was observed at both onset and response. **E**) Plots of the peak EEG frequency for each comparison frequency at onset and response; a null effect would lie along the dashed line. Shaded regions represent within-subject standard error.

**Figure 4:**
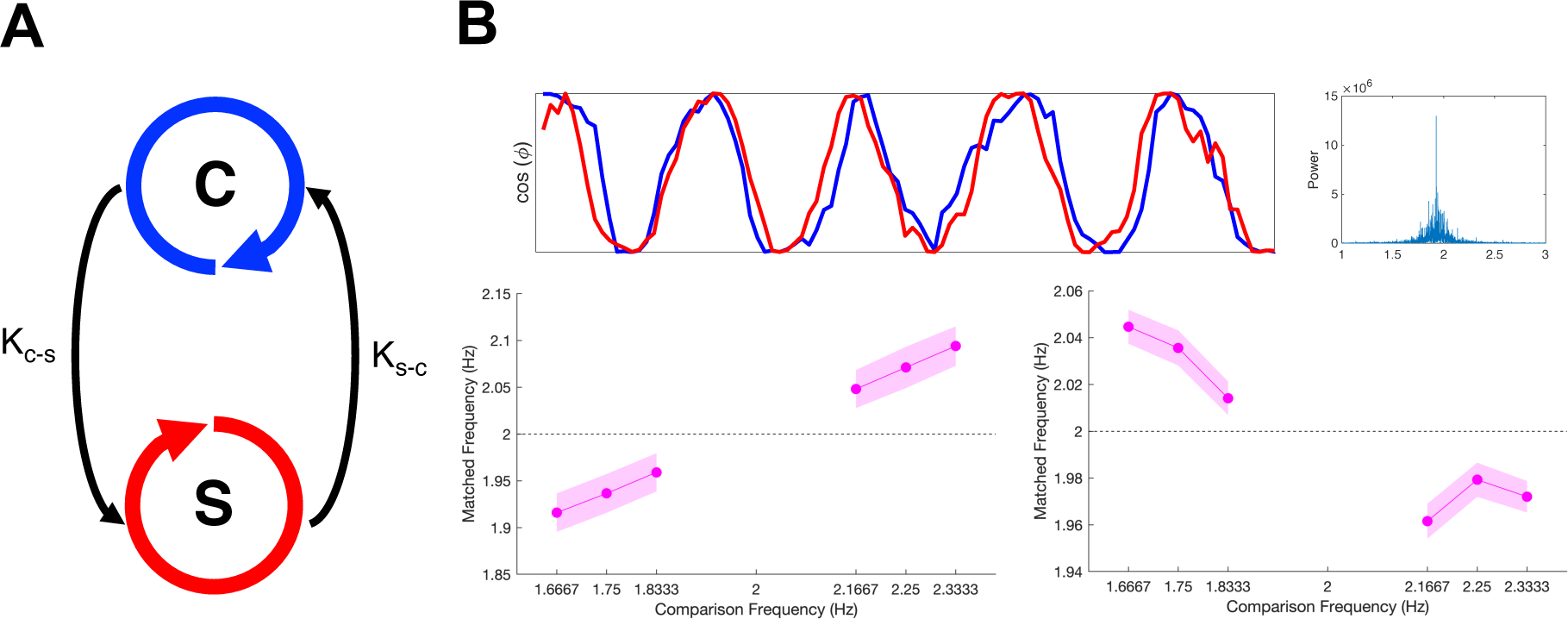
Coupled Oscillator Model. **A)** A simple model of two coupled oscillators for the standard (S) and comparison (C) rhythms, which were each externally driven by those rhythms. There is a coupling strength between both oscillators that “pull” them towards phase alignment with the parameters (K_S-C_ = 8, K_C-S_ = 3) such that the standard exerted a larger coupling strength upon the comparison. **B)** Each oscillator was corrupted by Gaussian noise, with no time-delay between oscillators, and run for 1000 seconds to achieve an equilibrium in coupling (Zalta, et al. 2020). On a given randomized “trial”, the oscillators were presented with the standard and comparison rhythm. We calculated the peak frequency of the standard and comparison oscillators via a fast Fourier transform as well as calculated the half-difference between the standard oscillator frequency on the previous and present trial (on every trial except the first) which was added to the standard oscillator’s initial frequency for the next trial. In this way, the standard oscillator shifted away from 2Hz by the amount that it had been pulled by the comparison frequency on the previous trial. We then calculated the average peak frequency of the comparison oscillator as a function of the previous trial’s comparison frequency. Bottom panels represent the average final frequency of C as a function of presented frequency on the present and previous trial replicating behavioral effects. Shaded regions represent standard error across 1000 simulations of 120 trials each.

### Experiment 2

If the bias and entrainment effects observed in behavior and EEG are the result of a system of coupled oscillators, a prediction then is that by altering the coupling strength of the oscillators, one should observe a resultant change in behavior. This question gets at the heart of dynamic theories of entrainment, as if entrained oscillators truly reflect the perception of rhythm then by altering their properties one should change perception. To test this, we applied 2Hz tACS over the SMA to a new group of participants (see Materials and Methods) performing this task. Stimulation was administered phase-locked to the start of the trial (and so the standard tempo), with true or sham stimulation in separate blocks. We additionally collected confidence judgments at the end of each trial to measure subjectively perceived performance.

Once again, we observed an effect of the comparison frequency such that responses were biased towards them on the present trial [*F*_(5,120_ = 33.087, *p*<0.001, η^2^*_p_* = 0.58] and away from them on the previous trial, although only at a trend level [*F*_(5,120_ = 2.059, *p*=0.075, η^2^*_p_* = 0.079]. Crucially, we observed an effect of stimulation compared to sham [*F*_(1,24)_ = 5.25, *p* = .031, η^2^*_p_* = 0.18; Fig. 5], such that participants showed significantly improved accuracy during the stimulation condition compared to sham in a linear fashion [*t*_(24)_ = −2.29, *p* = .031]. Further, we observed that stimulation also shifted the influence of the previous trial (*n* – 1), such that the stimulation condition was less influenced by the previous trial [*F*_(1,24)_ = 4.65, *p* = .041, η^2^*_p_* = 0.16; Fig. 5]. Further, the stimulation condition showed a linear shift away from the previous trial, closer to the target frequency [*t*_(110)_ = −2.16, *p* = .041].

**Figure 5:**
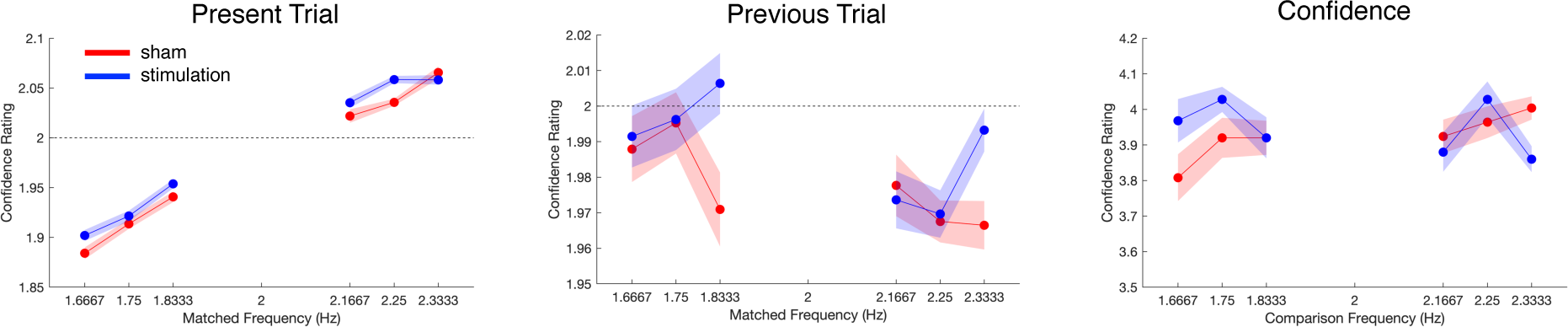
tACS results from Experiment 2. On the present trials, subjects again exhibited biases, yet were significantly attenuated following 2Hz tACS stimulation, such that responses were closer to the 2Hz standard tempo; notably, this effect was observed most strongly for slower comparison tempos. For responses based on the previous trial, stimulation also significantly reduced the bias exerted by the previous trial. Right panel displays confidence judgments, with no significant differences between stimulation conditions.

Finally, we investigated participants’ confidence ratings of how well they believed they corrected matched the comparison frequency to the target frequency. For this, we found no main effect for condition (*F*_(1,24)_ = .25, *p* = .62, η^2^*_p_* = 0.01; Fig. 5) or comparison frequency (*F*_(5,120)_ = .75, *p* = .59, η^2^*_p_* = 0.03; Fig. 5). As such, participants were not biased as to when they received stimulation compared to sham.

## Discussion

The data from Experiment 2 support our prediction from the behaviors exhibited in Experiment 1: rhythm perception was significantly altered when the coupling strength of the oscillators, in particular, the standard tempo, was changed. When participants underwent phase-locked stimulation, their accuracy was significantly improved compared to that of sham trials (Fig. 3A). Notably, application of stimulation resulted in a loss of bias to anchor to the trial’s comparison frequency and, indeed, pushed this bias effect in the opposite direction, resulting in significantly improved accuracy. These findings accord to the coupled oscillator model predictions (Fig. 4) such that the standard oscillator has a stronger “pull” when a second oscillator is introduced via stimulation.

In terms of our model, stimulation to the SMA via tACS may be increasing the coupling strength of K_S-C_, explaining the significant shift in accuracy towards the target frequency (Fig. 5). Further, when considering the effects of the previous trial on the current trial, our data from Experiment 2 more closely resemble the model for *n* – 1 compared to data from Experiment 1, suggesting that tACS is able to relieve some of this carryover bias.

### Entrainment

In line with previous studies, our data support entrainment primarily through increased accuracy during phase-locked stimulation in Experiment 2 (Fig. 5). As demonstrated by our data, phase-locked tACS delivered to the SMA bilaterally resulted in better performance and reduced anchoring to the comparison frequency compared to the sham blocks. Further, even though accuracy improved with tACS, confidence ratings were not different between stimulation and sham blocks, suggesting that subjects were unaware of their change in behavior. Overall, tACS delivered to the SMA modulated behavioral activity by increasing the coupling strength of the target frequency oscillator, resulting in a stronger entrainment to the target frequency. Our data therefore not only support the role of entrained oscillators in rhythm perception but are also indicative of enhanced temporal predictions as shown through improvements in accuracy.

Entrainment is also suggested in Experiment 1 by the presence of the alignment of peaks in the EEG recordings for trial onset to the target frequency (2Hz) as well as comparison frequencies (1.66 – 2.33 Hz). This is furthered by the shifting of peaks towards the target frequency and the disappearance of peaks at the comparison frequencies for trial offset (Fig. 2). The EEG data in Experiment 1 are congruent with previous studies in which the frontocentral region exhibited neural firing locking to external auditory stimuli (Cadena-Valencia et al., 2018; Fuijioka et al., 2012; Nozaradan et al., 2011). Yet, neural entrainment has been difficult to validate methodologically, primary given that its exact nature is still debated in the literature. One criticism of methodological evidence for entrainment is whether entrainment is a reflection of external stimuli in the auditory pathway or truly an endogenous representation of that stimuli (Breska & Deouell, 2017). As such, though phase-locking is an aspect of temporal predictions in general, it should not be overlooked when investigating entrainment, especially in regard to our data which showed that phase-locked stimulation significantly improved accuracy in the novel tempo matching task. Crucially, our data suggest that strengthening phase-locking of the SMA to exogenous rhythmic stimuli can improve temporal predictions.

Rosso and colleagues (2023) claimed that methodological evidence for entrainment relies on the use of perceptual and sensorimotor systems. Morillon and Baillet (2017) demonstrate that the motor system provides temporal context that is used for sensory processing as shown through temporal predictions being encoded by delta-beta oscillations in motor regions that are functionally connected with sensory regions. This inclusion of sensorimotor systems is supported by the necessity for predictions, an imperative component of entrainment which includes sensory and motor expectations based on a sensorimotor integration allowing for real time corrections when compared to sensory feedback, allowing for coupling of separate oscillatory systems (Ross & Balasubramaniam, 2014; Ross & Balasubramaniam, 2022). In addition to coupling, entrainment involves neural firing becoming frequency-and phase-locked (Haegens and Zion Golumbic, 2018; Lakatos et al., 2019; Ross & Balasubramaniam, 2022; Rosso et al., 2023). Not only has entrainment been shown to occur in both auditory and visual rhythmic paradigms, but tapping along to the exogenous beat has shown increased power and phase-coherence in EEG recordings (Comstock & Balasubramaniam, 2022). Indeed, entrainment involves alignment of not only neural firing but of internal motor and sensory rhythms themselves (Rosso et al., 2023).

It is suggested that the SMA engages in entrainment by acting on a dynamic motor plan that occurs before the completion of an interval to create an endogenous metronome (Cadena-Valencia et al., 2018; Cannon & Patel, 2021). Notably, the level of neural entrainment may be dependent on the presentation of the exogenous rhythm. Stupacher and colleagues (2016) found that entrainment was increased for drum rhythms compared to an isochronous metronome suggesting that entrainment involves endogenous timing processes rather than only physical input. As such, it is possible that entrainment is needed to help discover the beat when accompanied by overlapping musical stimuli and is not as prominent for beat maintenance alone.

That being said, it has been suggested that entrainment may enhance beat perception. The resonance theory for beat and meter processing postulates that the presence of neural entrainment resonating at the beat frequency acts as a metaphorical bridge for beat perception (Nozaradan et al., 2011). Additionally, this theory also states that an internal representation of the meter is supported by subharmonics of the external beat frequency (Nozaradan et al., 2011). As such, it is suggested that the SMA may act as an internal metronome to facilitate beat perception (Cadena-Valencia et al., 2018; Cannon & Patel, 2021). As demonstrated by our oscillatory model, this activity may be shifted depending on changes to the external auditory stimuli, supporting the dynamic role of the SMA in temporal prediction of external rhythmic stimuli.

### Conclusion

Overall, our findings support the theory of entrainment in the SMA, as shown by improved accuracy during stimulation trials in which tACS was frequency-and phase-locked to the target auditory beat. This is theorized to be accredited to the increased coupling strength that the stimulation provides to the standard tempo, as suggested by our adapted model. This work opens potential future avenues to explore and manipulate entrainment mechanisms in other paradigms, as well as in pathologies where dysfunction of rhythmic processing is evident (Breska & Ivry, 2018; Nozaradan, et al. 2017).

## Materials and Methods

### Experiment 1

#### Participants

For this study, 25 healthy participants (9 females, 16 males) ages 18-36 years old (*M* = 24.64 years old) from on or around George Mason University volunteered and were compensated for their time. Two participants were dropped due to incomplete data resulting in the use of 23 participants for data analysis. Each participant was right-handed and had normal or normal-to-corrected vision. Of the 22 responses, 16 participants reported having formal musical training and/or play an instrument (M = 7.53) with an average of 5.93 years since they last played. The experiment was approved by George Mason University’s Institutional Review Board before data collection began.

#### Materials

PsychoPy (v.2021.2.3) was used to create the tempo matching task. The auditory stimulus for the bass tone was generated in GarageBand while the musical tone (A4 at 440 Hz) was generated in Audacity (v.3.2.5). Both stimuli were then altered using Audacity to match for duration, loudness, and fading. Participants were seated in front of a 32” LCD monitor (Display++, Cambridge Research) running at 120Hz refresh rate with a speaker on both sides of the monitor. A consistent volume was maintained across participants.

#### Procedure

Participants were tested on George Mason University’s campus. Upon arrival to the labspace, participants gave informed consent and completed a demographic questionnaire, the Edinburgh Handedness Inventory, and a musical experience form. Each participant underwent EEG which consisted of using 64-actiCAP Slim electrodes connected to an actiCHamp amplifier (Brain Products). EEG data was collected via BrainVision Recorder software.

The experiment began by collecting baseline data in which participants listened to 10 trials of only the bass tone totaling 5 minutes followed by three blocks of 60 trials of a tempo matching task. For the tempo matching task, participants were presented with two simultaneous tones (a bass tone and a musical tone) that were repeated at different frequencies dependent on trial. The bass tone maintained a steady frequency of 2 Hz across trials while the musical tone was played at either 1.66, 1.75, 1.83, 2.16, 2.25, or 2.33 Hz.

For each trial, participants were asked to modulate the frequency of the musical tone (the comparison frequency) to match that of the bass tone (the target frequency) through the use of the up and down keys on the keyboard which changed the comparison frequency by .02 Hz in the desired direction. Once they deemed that they had successfully matched the comparison frequency to that of the target, the participant pressed the spacebar to continue on to the next trial.

#### EEG Data Collection and Preprocessing

EEG activity was sampled at 1000 Hz. Offline data analysis was conducted using EEGLAB. The mastoid (TP9 and TP10) and temporal (FT9 and FT10) channels were removed for noise and the data were re-referenced to common average. The data were then epoched for the onset and offset of the trial for each condition as well as up/down button presses. For the onset of the trial, the epoch length extended from −1s to 10s, with a baseline correction of −1s to 0. For the offset of the trial, the epoch length extended from −10s to 1s relative to the button-press used to terminate the trial; data were mean-centered, rather than baseline-subtracted. All data were then bandpass filtered between 0.01 and 50Hz using the EEGLAB function *pop_eegfiltnew.* Independent Component Analysis (ICA) was then run for each epoch type for manual removal of noisy components due to eye and muscle movements.

#### Behavioral Data Analysis

Behavioral responses were analyzed by calculating the frequency at the end of each trial, when subjects pressed the space bar to indicate that the comparison and standard frequencies were matched. The mean frequency was calculated for each of the comparison frequencies. Additionally, we calculated the mean trial time and number of up/down button-presses for each comparison frequency, as well as the mean time from the onset of the trial to when subjects first began adjusting the comparison frequency and the mean time from when subjects made their final adjustment to when they pressed the space bar. We also calculated the average frequency by the comparison frequency presented on the previous trial. All behavioral data were analyzed using JASP (https://jasp-stats.org).

#### EEG Data Analysis

EEG data were analyzed by taking the timeseries data for onset and offset-locked epochs and converting them to frequency representations using a fast Fourier transform as implemented in EEGLAB using the *spectopo* function with a frequency resolution of 0.091Hz. Frequency data were then detrended with a 2^nd^ order polynomial using Matlab’s *detrend* function. For both onset and offset data, we calculated the average normalized power for trials presented at each comparison frequency for each of the possible comparison frequencies. For example, on trials where the comparison frequency was 1.66Hz, we calculated the power at each frequency band of the *possible* comparison frequencies (1.66 – 2.33Hz), yielding a 6×6 matrix. From these values, for each comparison frequency trial, we measured the peak frequency as the band with the highest power. These values were then analyzed via a repeated-measures ANOVA with comparison frequency as a within-subjects factor.

### Experiment 2

#### Participants

For this study, 25 healthy participants (10 females, 15 males) ages 18-33 years old (*M* = 20.71 years old) from on or around George Mason University volunteered and were compensated for their time. Each participant was right-handed and had normal or normal-to-corrected vision. Six participants reported having formal musical training and/or play an instrument (M = 6.92). The experiment was approved by George Mason University’s Institutional Review Board before data collection began.

#### Materials

In addition to the same tempo matching task from Experiment 1, participants were asked to rate their confidence for each trial using a slider. Participants were seated in front of a 24” Dell S2417DG monitor running at a 100Hz refresh rate with a speaker on both sides of the monitor. A consistent volume was maintained across participants.

#### Procedure

Participants were tested on George Mason University’s campus. Upon arrival to the lab space, participants gave informed consent and completed a demographic questionnaire, the Edinburgh Handedness Inventory, and a musical experience form. Each participant underwent tACS using the neuroConn DC-Stimulator which was placed over the SMA bilaterally (FC1 and FC2) using the 10-20 system; this montage has been previously used by our group, with source modeling demonstrating maximal current induced at the SMA (Wiener, et al. 2018). The neuroConn DC-Stimulator was set to 1mA with a frequency of 2 Hz and a phase of 90° such that it matched the attack and decay of the target tone stimuli. In this way, the stimulator was set to be frequency-and phase-locked to the target tempo.

The experiment consisted of four blocks of 30 trials. Stimulation blocks consisted of stimulation being delivered from the onset of the tempo matching task and ended once the participant pressed the spacebar. Stimulation was phase-locked to the target frequency (2 Hz). Sham blocks consisted of stimulation being delivered for one second at the onset of each tempo matching task. The experiment consisted of a within-subjects design in which each participant received alternating stimulation and sham blocks which was counter-balanced across participants. Subjects completed a poststudy questionnaire to assess side effects (tingling, itching sensation, burning sensation, pain, and fatigue). All participants were actively encouraged to report any perception of tACS-induced phosphenes throughout the experimental sessions.

#### Behavioral Data Analysis

The same behavioral data analyses from Experiment 1 were run for Experiment 2. In addition, we also calculated the average confidence rating for each comparison frequency. All behavioral data were analyzed using JASP (https://jasp-stats.org).

#### Computational Modeling

A model of coupled oscillators (Fig. 4) was constructed based on previous work (Zalta et al., 2020; Petkoski et al., 2018; Kuramoto, 1975). For this approach, we assumed that two internal oscillators existed for the standard (S) and comparison (C) rhythms, which were each externally driven by those rhythms. The model assumed a coupling strength (K) between each of the oscillators with one another, such that the standard rhythm “pulled” the comparison towards phase alignment with it K_S-C_, while comparison also exerted its own coupling strength K_C-S_ with the parameters (K_S-C_ = 8, K_C-S_ = 3) such that the standard exerted a larger coupling strength upon the comparison, but the comparison still had some influence on the standard. Each oscillator was additionally corrupted by Gaussian noise, with no time-delay between oscillators, and run for 1000 seconds to achieve an equilibrium in coupling (Zalta, et al. 2020).

To apply the model to our data, we took all six possible comparison rhythms and generated a randomized trial list of the same length as our human participants. On a given “trial”, the oscillators were presented with the standard and comparison rhythm; after this, we calculated the peak frequency of the standard and comparison oscillators via a fast Fourier transform. The frequency of the comparison oscillator was thus used as the model’s “choice” for that trial; the average of each choice was calculated to determine model performance across the six tested comparison rhythms. To account for the repulsive bias, on every trial beyond the first, we calculated the half-difference between the standard oscillator frequency on the previous and present trial; this amount was then added to the standard oscillator’s initial frequency for the next trial. In this way, the standard oscillator shifted away from 2Hz by the amount that it had been pulled by the comparison frequency on the previous trial. We then calculated the average peak frequency of the comparison oscillator as a function of the previous trial’s comparison frequency, as done for the behavioral data.

